# Towards the development of an insulin degradation test

**DOI:** 10.64898/2026.02.12.705529

**Authors:** David Ritz, Elizabeth-Lauren Stevenson, Daniel Schultz

**Affiliations:** Department of Microbiology & Immunology – Geisel School of Medicine at Dartmouth,Hanover, NH 03755, USA; Department of Molecular and Systems Biology – Geisel School of Medicine at Dartmouth, Hanover, NH 03755, USA

**Keywords:** insulin, fibrillation, degradation, sensing

## Abstract

People with diabetes rely on exogenous insulin to reduce blood glucose levels, compensating for insulin resistance or impaired pancreatic β-cell function. Despite being essential for diabetes management, insulin formulations exhibit inconsistent performance due to their relatively fragile stability. This instability carries significant cost implications: some individuals spend over $1,000 USD per month on insulin, and these high prices influence one in six Americans with diabetes to ration their insulin supplies. Environmental stressors can induce conformational changes that cause insulin to misfold and aggregate into fibrils, which are inactive structures that contribute to long-term diabetic complications. Although insulin’s instability is well-documented, no test currently exists outside of laboratory settings to determine whether an insulin formulation has degraded. Here, we compare biochemical techniques for assessing bioactivity and structural integrity in three commercial insulin analogs exposed to physiologically relevant stress conditions, showing that fibril formation precedes measurable loss of bioactivity in insulin and that fibrillation depends on both the stressor type and the insulin formulation tested. We then demonstrate proof-of-concept testing for antibody-based degradation detection using commercial monoclonal antibody candidates. Together, these findings underscore the critical need for accessible insulin quality testing and demonstrate the feasibility of antibody-based detection of insulin fibrillation.

**Importance:** Insulin remains one of the most essential yet fragile biopharmaceuticals used in modern medicine. Globally, 150 million people with diabetes depend on exogenous insulin to regulate blood glucose levels, but the protein’s inherent instability can cause degradation during storage or transport. These degradation events can reduce insulin’s potency and safety, yet patients and healthcare providers currently have no practical means to assess insulin quality before injection. This knowledge gap contributes to inconsistent therapeutic outcomes and increases the risk of complications associated with degraded insulin. Our work directly addresses this unmet clinical and public health need by identifying the molecular changes that occur when insulin analogs are exposed to everyday environmental stressors and by completing proof-of-concept testing for a device to detect insulin fibrillation. We demonstrate that structural transitions to fibrils precede measurable loss of bioactivity and that fibrillation behavior depends on both the insulin analog and the stressor type. By combining biochemical characterization with antibody-based detection, this study establishes a foundation for a low-cost, accessible method to verify insulin integrity outside the laboratory. Such a tool could prevent the use of degraded insulin, improve treatment consistency, and empower patients to ensure the quality of their medication. More broadly, this approach exemplifies how protein stability monitoring can be integrated into biotherapeutic quality assurance, improving safety, efficacy, and trust in life-sustaining biologic medicines.

## Introduction

Insulin manufacturers sell analog insulin to diabetics that differs by a few amino acid substitutions relative to human insulin, changing insulin’s onset of action^1^. Although an essential medication for a person with diabetes (PwD), human and analog insulin are well-documented to have an inconsistent effect due to their relatively fragile stability^2–4^. Insulin can be degraded by environmental factors during storage, such as excessive heat, agitation, or other stressors, despite the preservatives that are added in analog insulin formulations^5^. When insulin is challenged by environmental stressors, it undergoes conformational changes that initiate fibrillation – a multistep misfolding pathway in which native α-helical insulin forms partially unfolded, aggregation-prone fibril intermediates that can assemble into mature β-sheet–rich amyloid fibrils^6^. Progression along the fibrillation pathway is associated with loss of native α-helical structure and increasing β-sheet content, changes that impair insulin’s ability to bind and activate the insulin receptor^7–11^. The presence of fibrils in an insulin solution can lead to catheter occlusion^12^ and reduced bioactivity^13^, which can cause hyperglycemia and ketoacidosis for diabetics^14,15^.

Compounding this problem, persons with diabetes are often unaware when their insulin has been compromised, creating a dangerous clinical dilemma^4,15^. A PwD may administer insulin without observing the expected reduction in blood glucose levels, forcing them to decide whether to increase their dosage – risking hypoglycemia if the insulin is functional – or to wait longer for the insulin to take action – risking hyperglycemia and ketoacidosis if the insulin is impaired. In this situation, the PwD must also decide whether to keep using the insulin of unknown bioactivity for future administrations or to dispose of the insulin supply. This is a costly choice, as some persons with diabetes spend over $1000 USD on insulin per month^16^. These high prices already influence 1 in 6 Americans with diabetes to ration their insulin and use it past strict expiration dates written on the medication’s packaging^17^. Further, chronic injection of fibril-containing insulin can lead to insulin-derived amyloidosis^18^, which is described as a subcutaneous degraded insulin mass that forms due to frequent insulin injections at an injection site, leading to a sustained immune response at the lesion^19^ and unpredictable glycemic control^20^. Persons with diabetes are especially susceptible to insulin-derived amyloidosis due to a need for frequent insulin injections over their lifetime and a limited number of injection sites on their bodies^19^.

Despite the demonstrated fragility of insulin and clinical need for degradation detection before injection, no test currently exists outside of a lab setting to determine if insulin has become degraded. Here, we use multiple biochemical techniques for assessing bioactivity and structural integrity in three commercial insulin analogs exposed to physiologically relevant stress conditions, showing that fibrillation precedes measurable loss of bioactivity in insulin and that degradation depends on both the stressor type and the insulin formulation tested. Given that recent studies have shown that conclusions about insulin’s fidelity are method dependent^21–25^, we applied several complementary analytical approaches to characterize degradation. Finally, we demonstrate antibody testing capable of detecting thermally-formed insulin fibrils using commercial monoclonal antibody candidates, which could be incorporated into a future point-of-care assay for a PwD. Together, these results underscore the need for accessible insulin quality testing and demonstrate the feasibility of antibody-based detection of insulin degradation.

## Results

### Insulin cloudiness is not indicative of fibrillation

Insulin manufacturers recommend not administering insulin that has become cloudy^26–28^, as this can be an indication of a degradation product precipitating or a sign of contamination. To determine whether this user visual inspection test can detect insulin fibrils, we first degraded insulin using a range of relevant stressors for up to 96 hours to assess whether the solutions became cloudy in their original formulations. We tested three widely used insulin brands: Humalog (fast-acting), Novolog (fast-acting), and Basaglar (long-acting)^29^. Each insulin was separately exposed to -20°C, 37°C, 37°C with agitation, 65°C, air, agitation, UV light, and ideal conditions at 4°C, spanning environmental conditions that an insulin vial may encounter during its lifetime.

Cloudiness was first quantified using a spectrophotometric absorbance assay, which measures light scattering from insoluble aggregates (**Fig. 1A**). Across all insulins and conditions, only Humalog incubated at 65°C exceeded an absorbance of 0.1 and showed a detectable increase in cloudiness by eye (**Fig. S1**). To assess fibril formation during these stress trials, we next used a Thioflavin T (ThT) fluorescence assay (**Fig. 1B, Fig. S2-S3**). ThT brightly fluoresces after binding to fibril species, recognizes fibril structures in seconds^30^, and is believed to have an affinity to general epitopes of mature insulin fibrils and their intermediates^31–33^. Humalog incubated at 65°C caused high levels of fibrillation, agreeing with a high absorbance reading. In contrast, Novolog incubated at 65°C and Basaglar incubated at 37°C with agitation also developed high levels of fibrillation but exhibited little changes in absorbance, making these degraded samples visually indistinguishable from non-degraded insulin (**Fig. S1**).

**Figure 1:**
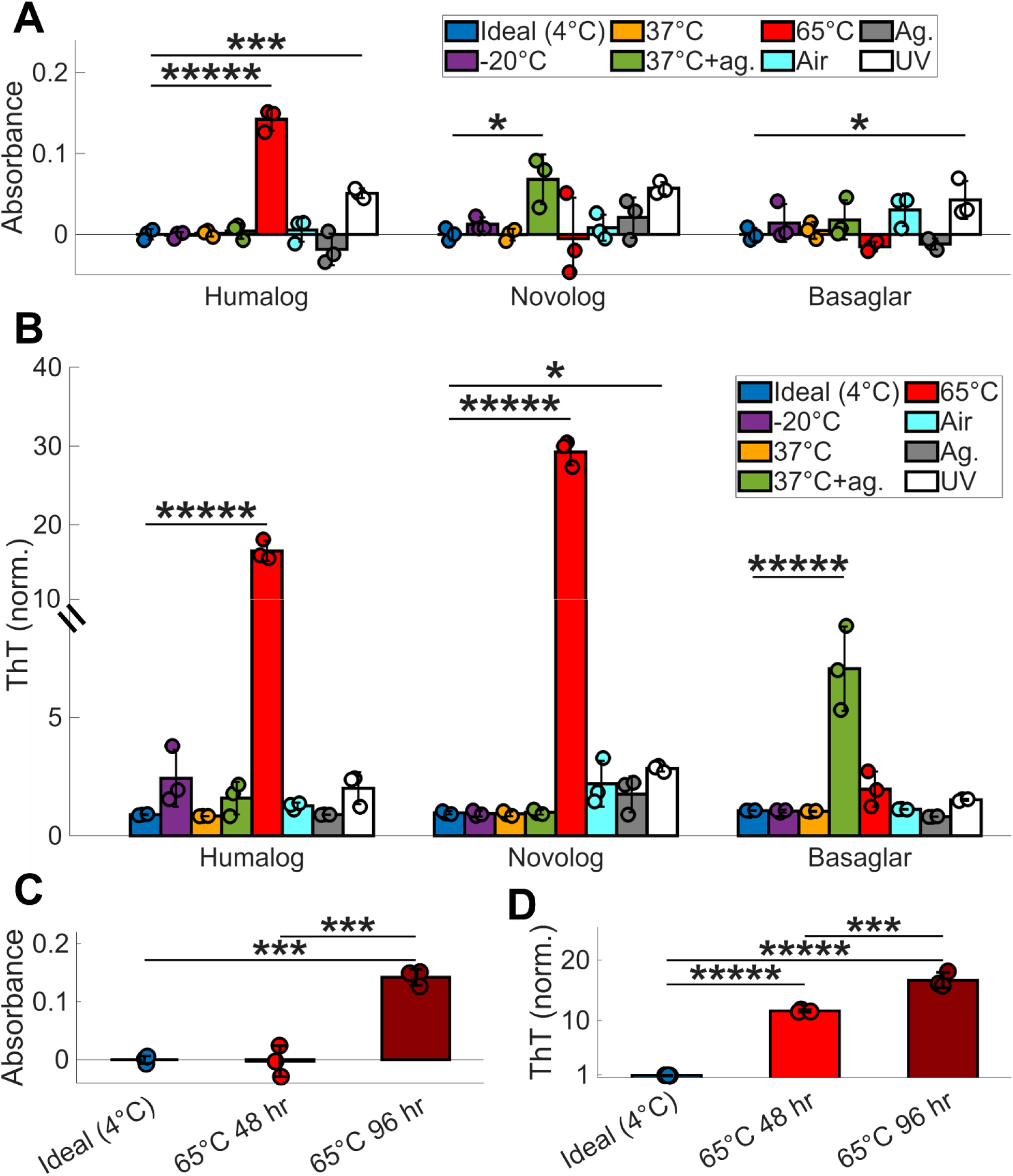
Insulin absorbance readings do not detect insulin degradation. **(A)** Absorbance at 600 nm and **(B)** ThT fluorescence of Humalog, Novolog, and Basaglar after 96 hours of exposure to each environmental stressor, except for the 302 nm UV light trial, which was exposed for only 2 hours due to rapid deterioration and color change of the insulin after this point. 3 replicates were used for each trial, and all trials were compared to 4°C using ANOVA with Dunnett correction. **(C)** Absorbance at 600 nm and **(D)** ThT fluorescence of unaltered Humalog insulin at ideal storage conditions of 4°C, Humalog at 65°C for 48 hours, and Humalog at 65°C for 96 hours. 3 replicates were used for each trial, and each pairwise comparison was assessed using ANOVA with Tukey-Kramer correction. The asterisks denote ***** p < 10^-5^, **** p < 10^-4^, *** p < 10^-3^, ** p < 10^-2^, and * p < 5*10^-2^.

We next examined whether fibril formation precedes visible cloudiness by following the kinetics of Humalog degradation at 65°C. Although an increase in absorbance was detectable only at 96 hours of incubation (**Fig. 1C**), substantial fibril formation was already present by 48 hours and increased further by 96 hours (**Fig. 1D**). Thus, a cloudiness check by a Humalog insulin user would identify degradation only at a late stage, well after significant fibrillation had already accumulated. Together, these results show that user visual inspection tests recommended by insulin manufacturers fail to reliably detect fibrillation across stress conditions and insulin formulations.

### Insulin fibrillation precedes measurable loss of bioactivity

There are two clinically relevant consequences of insulin fibrillation: (1) formation of fibrillar aggregates, which can contribute to long-term diabetic complications^18^, and (2) loss of functional insulin, which reduces potency. Importantly, the thresholds at which these two effects become clinically meaningful may not coincide, and the extent to which fibril formation compromises insulin bioactivity remains unclear. To investigate this, we generated insulin samples spanning a range of fibrillation states and evaluated their function using a cell-based insulin bioactivity assay described by the FDA^34^. In this assay, diluted insulin samples are applied to Chinese Hamster Ovary (CHO) cells expressing human insulin receptor B, and bioactivity is quantified by measuring insulin-induced receptor activation. To induce fibrillation, each insulin analog was exposed to environmentally relevant stressors, including heat, agitation, freeze–thaw cycling, and expired storage. These treatments generated diverse fibrillation profiles across Humalog, Novolog, and Basaglar, enabling the comparison between the concentration of fibril species and loss of bioactivity (**Fig. 2**). Bioactivity was quantified from the area under dose–response curves generated from multiple dilutions of each degraded insulin sample, allowing functional changes to be assessed across a physiologically relevant concentration range (**Fig. S5**; **Methods**).

**Figure 2:**
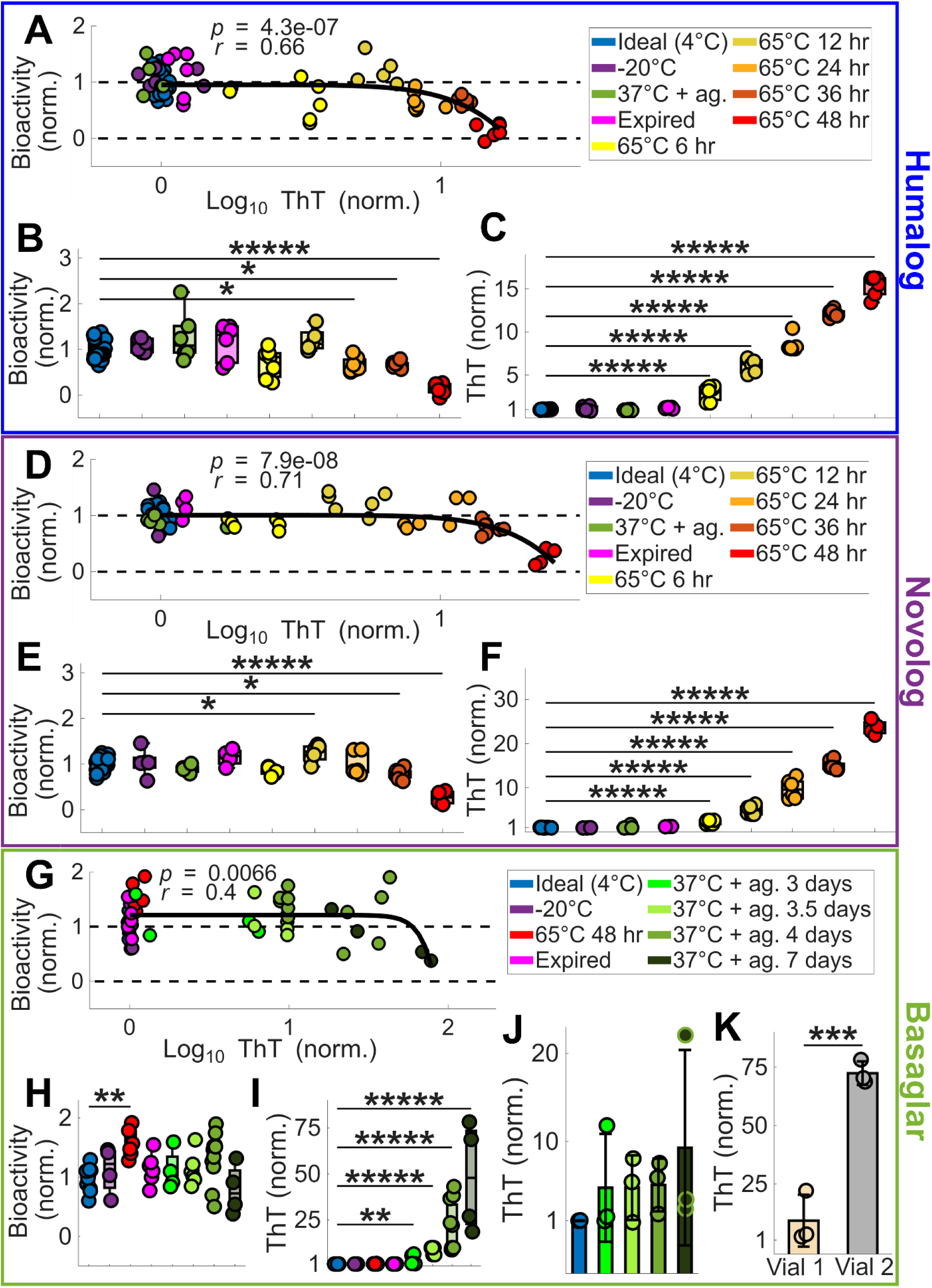
Insulin’s bioactivity decreases when degradation is above analog-specific thresholds. **(A)** CHO cell bioactivity plotted against ThT fluorescence for Humalog degraded by various methods. For each scatter plot in this Figure, the black curve is a power law function fit to the scatterplot data, with the *p*-value and Pearson’s *r* value of the fit displayed. For all CHO experiments, 4 to 27 replicates were used for each stress condition (**Supplemental Data 2**). **(B)** The CHO cell bioactivity and **(C)** ThT fluorescence of each degradation method for Humalog are shown. All trials in B and C were compared to 4°C using ANOVA with Dunnett correction. **(D)** CHO cell bioactivity plotted against ThT fluorescence for Novolog degraded by various methods. **(E)** The CHO cell bioactivity and **(F)** ThT fluorescence of each degradation method for Novolog are shown. All trials in E and F were compared to 4°C using ANOVA with Dunnett correction. **(G)** CHO cell bioactivity plotted against ThT fluorescence for Basaglar degraded by various methods. **(H)** The CHO cell bioactivity and **(I)** ThT fluorescence of each degradation method for Basaglar are shown. **(J)** Aliquots of one Basaglar vial were either kept at ideal conditions or were degraded at 37°C with agitation for 3, 3.5, 4, or 7 days. All trials in H, I, and J were compared to 4°C using ANOVA with Dunnett correction. **(K)** Two different vials of Basaglar were aliquotted and degraded at 37°C with agitation for 7 days. 3 aliquots were tested from each vial and were compared by a two-way t-test. The asterisks denote ***** p < 10^-5^, **** p < 10^-4^, *** p < 10^-3^, ** p < 10^-2^, and * p < 5*10^-2^.

Across all three insulin analogs, bioactivity remained largely intact until fibrillation surpassed an analog-specific threshold, revealed by fitting each dataset to a power-law function (**Fig. 2**). Humalog and Novolog had significant, strong correlations between fibril concentration and loss of bioactivity (Humalog: *p* < 10⁻^6^; *r* = 0.66; Novolog: *p* < 10⁻^7^; *r* = 0.71), while Basaglar had a significant, moderate correlation (*p* < 10⁻^2^; *r* = 0.40).

For Humalog, bioactivity decreased only after fibril levels reached ∼8-fold above fresh insulin (**Fig. 2A**), corresponding to 24 hours of incubation at 65°C (**Fig. 2B**). Yet fibrillation increased substantially after only 6 hours of incubation (**Fig. 2C**), demonstrating a delay between fibril formation and functional decline. For Novolog, bioactivity dropped once fibril levels reached ∼15-fold above fresh insulin (**Fig. 2D**), which corresponded to 36 hours at 65°C (**Fig. 2E**). As with Humalog, fibrillation rose sharply by 6 hours at this temperature (**Fig. 2F**), again indicating early structural degradation without immediate functional consequences. For Basaglar, bioactivity did not decline until fibril levels reached ∼70-fold above fresh insulin (**Fig. 2G**). Notably, none of the tested Basaglar groups showed a statistically significant loss of bioactivity (**Fig. 2H**), even though fibril concentrations increased markedly after 3 days at 37°C with agitation (**Fig. 2I**). Heterogeneity in Basaglar’s degradation likely contributed to the lack of bioactivity significance among experimental groups, as aliquots from the same insulin pen (**Fig. 2J**) or from different pens (**Fig. 2K**) showed different fibrillation fates under identical stress conditions.

Together, these results demonstrate that insulin analogs can accumulate substantial fibril loads without detectable loss of bioactivity in the CHO cell assay, but each analog exhibits a different threshold beyond which biological function becomes compromised.

### Insulin analogs decrease α-helix signatures when degraded

Although insulin is known to decrease its α-helix secondary structure after degradation^35^, it is unclear whether this transition occurs similarly across different insulin analogs. To investigate this, we prepared circular dichroism (CD) samples of Humalog, Novolog, and Basaglar by exposing each analog to its most fibril-inducing stressor: 65°C for Humalog and Novolog, and 37°C with agitation for Basaglar. These conditions generated insulin samples with undetectable, intermediate, and high fibril concentrations, spanning the range observed in the CHO bioactivity assays (**Fig. 3A**). Changes in secondary structure of the protein population were then assessed by CD, where reductions in the magnitude of the 208 nm and 222 nm minima indicate a loss of α-helical structure, a 222/208 nm ratio greater than one reflects a coiled-coil signature, and a singular minimum near 216 nm indicates β-sheet formation^36^. These spectral features enable direct comparison of structural changes among insulin analogs and correlation of those changes with fibril formation and bioactivity.

**Figure 3:**
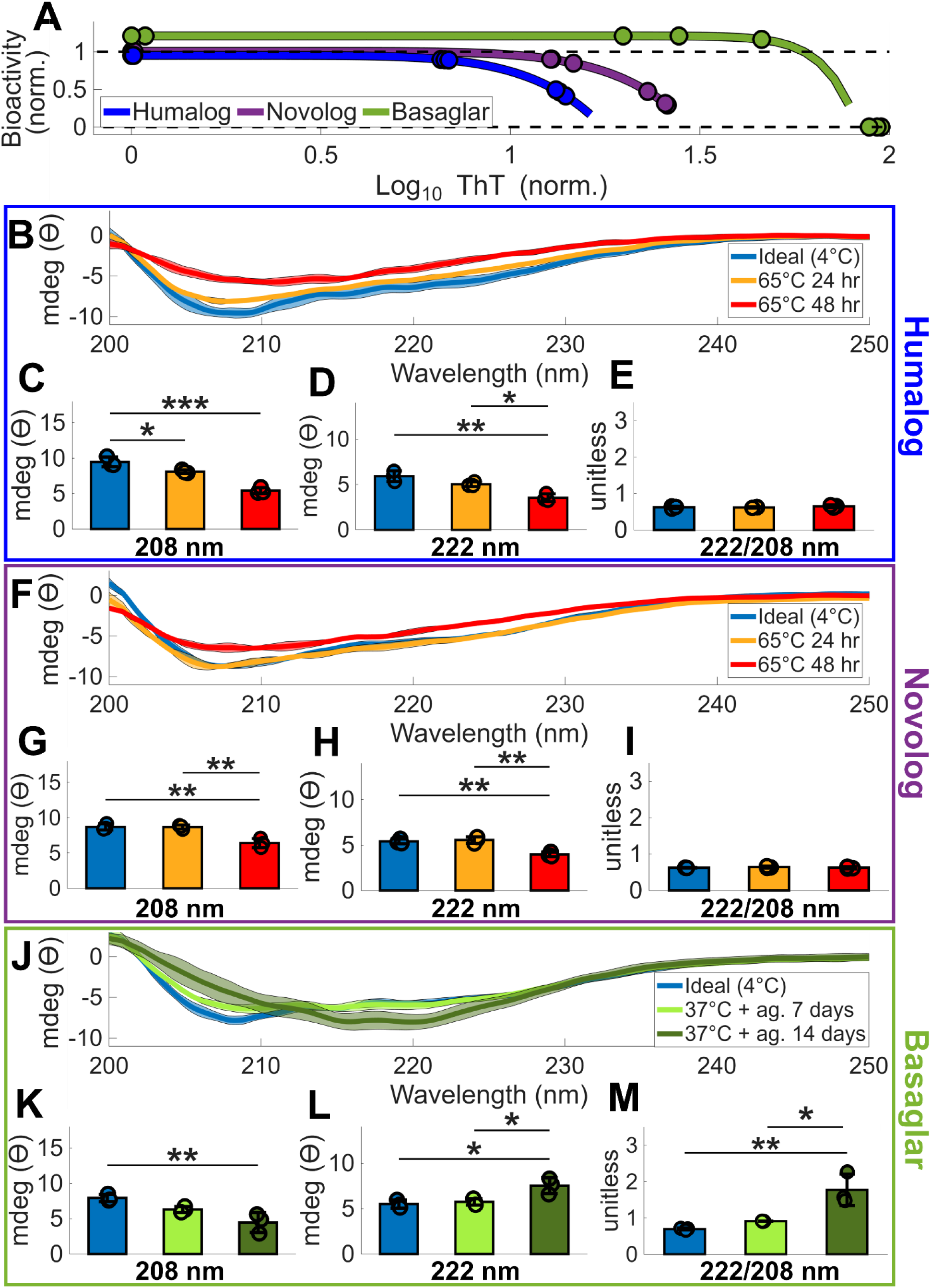
Insulin’s secondary structure decreases α-helix signatures during degradation. **(A)** Far-UV CD spectra were measured of degraded Humalog, Novolog, and Basaglar insulin samples that span the CHO bioactivity curves from Fig. 2. The dots indicate each CD sample’s ThT measurement and corresponding projected bioactivity. **(B)** CD spectra of Humalog insulin exposed to ideal storage conditions of 4°C, 65°C for 48 hours, and 65°C for 96 hours. For all spectra in this Figure, the bold line is the mean of three replicates, and the shading around each mean spectrum is ± the standard deviation. The magnitude of **(C)** 208 nm, **(D)** 222 nm, and **(E)** 222 nm divided by 208 nm CD measurements are quantified. These wavelength measurements are indicative of protein secondary structure. **(F)** CD spectra of Novolog insulin exposed to ideal storage conditions of 4°C, 65°C for 48 hours, and 65°C for 96 hours. The magnitude of **(G)** 208 nm, **(H)** 222 nm, and **(I)** 222 nm divided by 208 nm measurements are quantified. **(J)** CD spectra of Basaglar insulin exposed to ideal storage conditions of 4°C, 37°C and agitation for 7 days, and 37°C for 14 days. The magnitude of **(K)** 208 nm, **(L)** 222 nm, and **(M)** 222 nm divided by 208 nm measurements are quantified. In the bar graphs, each pairwise comparison was assessed using ANOVA with Tukey-Kramer correction. The asterisks denote ***** p < 10^-5^, **** p < 10^-4^, *** p < 10^-3^, ** p < 10^-2^, and * p < 5*10^-2^.

For Humalog, both 65°C incubation periods caused a decrease in the α-helix signature (**Fig. 3B**), with dampening of the characteristic minima at 208 nm (**Fig. 3C**) and 222 nm (**Fig. 3D**), evident after 24 hours of incubation. Neither incubation produced a coiled-coil signature (**Fig. 3E**). Despite the significant secondary-structure changes detected after 24 hours, degraded samples did not fall within the fibrillation range that reduces bioactivity (**Fig. 3A**), suggesting that a suitable fraction of Humalog still remains structurally intact at this degradation stage. The decrease in both the 208 and 222 nm minima without the emergence of a 216 nm band in these degraded samples indicates loss of native α-helical structure and formation of fibrillar intermediates rather than mature β-sheet fibrils^37^.

For Novolog, a reduced α-helix signature emerged only after the full 48-hour 65°C incubation (**Fig. 3F**). This condition dampened the minima at 208 nm (**Fig. 3G**) and 222 nm (**Fig. 3H**), consistent with reduced α-helices, but did not produce evidence of a coiled-coil structure (**Fig. 3I**). As with Humalog, the absence of a 216 nm minimum indicates formation of fibrillar intermediates rather than fully developed β-sheet fibrils in the degraded samples.

In contrast, Basaglar incubated for 14 days at 37°C with agitation developed distinct coiled-coil signatures (**Fig. 3J**). As the 208 nm trough dampened (**Fig. 3K**), the 222 nm trough’s magnitude increased (**Fig. 3L**), yielding an elevated 222 nm to 208 nm ratio characteristic of a coiled-coil signature^36^ (**Fig. 3M**). In addition, the emergence of a minimum near 216 nm in the samples degraded for 14 days is consistent with progression toward a β-sheet-rich mature fibril structure.

These data show that all three insulin analogs lose native α-helical structure and populate fibrillar conformations during degradation, with Basaglar uniquely progressing through a coiled-coil intermediate and showing the greatest shift toward β-sheet-rich organization.

### Thermally-degraded insulin products can be detected with monoclonal antibodies

The CD measurements suggest that degradation is associated with loss of native α-helical structure in all three insulin analogs, with accumulation of fibrillar intermediates and, in the case of Basaglar, β-sheet fibril conformations. Therefore, we next asked if degraded insulin can be detected using commercial monoclonal antibodies that recognize the secondary structure motifs of fibrillated proteins^38,39^. Such antibodies could form the basis of a simple point-of-use lateral-flow assay to determine whether insulin has fibrillated prior to injection. To evaluate antibody performance, we used dot blots to test the affinity of seven fibril-reactive antibodies towards degraded insulin analogs at undetectable, low, and high fibril concentrations. These degradation products were generated under several stress conditions to assess whether antibody binding is robust to the structurally distinct fibrils produced by distinct degradation routes (**Fig. 4A-B**).

**Figure 4:**
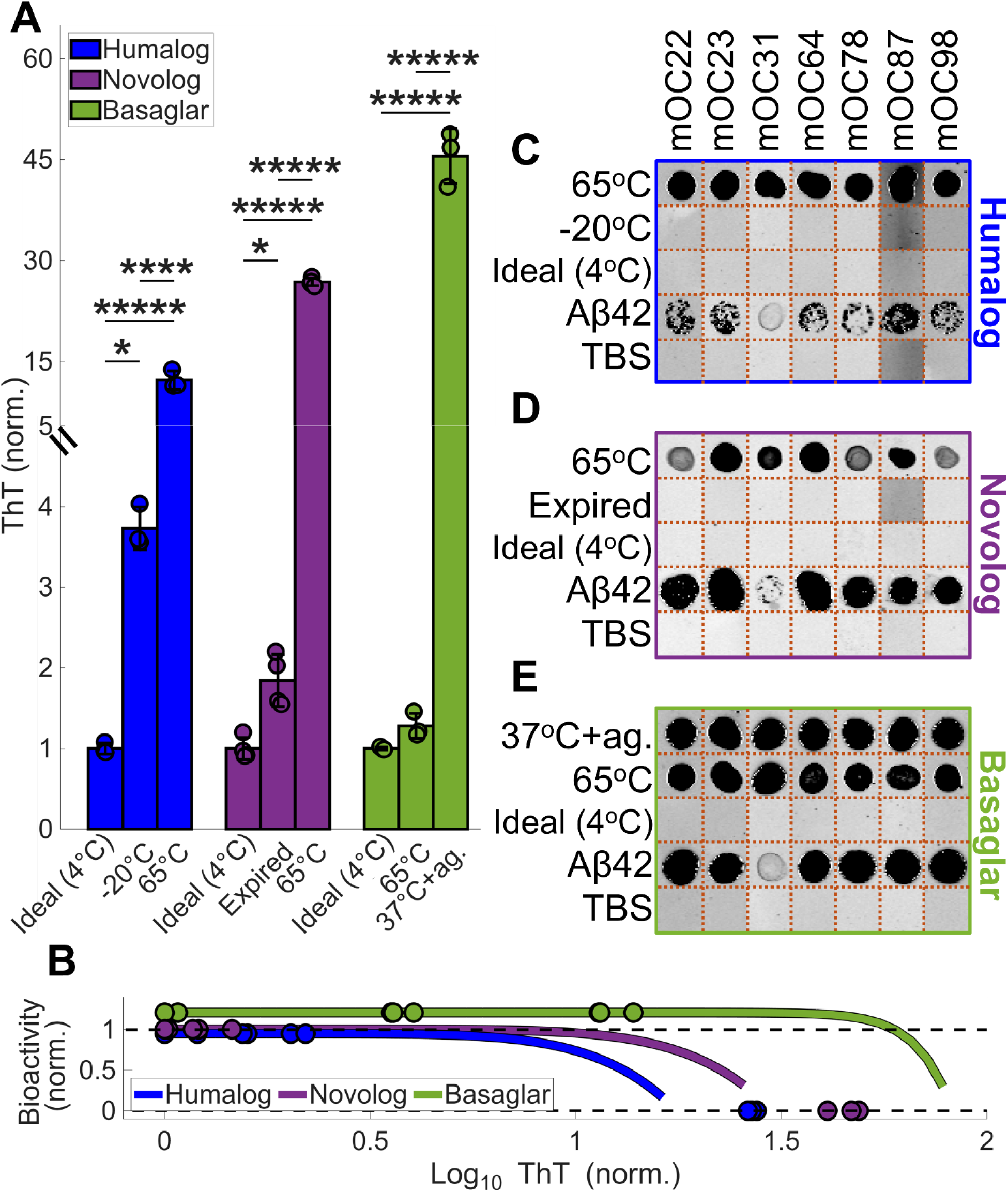
Antibody panel can detect fibril proteins formed from thermal exposures. **(A)** ThT readings of the insulin used for the dot blot experimentation. Each pairwise comparison was assessed using ANOVA with Tukey-Kramer correction. The asterisks denote ***** p < 10^-5^, **** p < 10^-4^, *** p < 10^-3^, ** p < 10^-2^, and * p < 5*10^-2^. **(B)** The ThT readings of the insulin analogs are denoted as dots on the CHO bioactivity curves from Fig. 2, showing the samples’ projected bioactivities. Results of the fibril antibodies’ dot blots for **(C)** Humalog, **(D)** Novolog, and **(E)** Basaglar are shown in compiled images. Each antibody is denoted a different mOC number. Aβ42 was the positive control, and TBS was the negative control. Each antibody per insulin type was a separate experiment and was cropped together to create panels **C-E**. The orange dotted lines are where the images were cropped. The full dot blot membranes are shown in **Fig. S8**.

While no antibodies had detectable binding to the fresh Humalog control, all antibodies bound to Humalog incubated at 65°C for 48 hours (**Fig. 4C**). Notably, there was no antibody binding to freeze-thaw cycled Humalog, even though these samples had detectable fibrillation during the ThT assay. This suggests that the antibodies have different affinities to different fibril conformations, and that the antibodies and ThT may recognize different motifs.

A similar pattern was observed for Novolog. Like Humalog, Novolog had no antibody recognition of the fresh insulin control. Further, while all antibodies bound strongly to Novolog incubated at 65°C for 48 hours (**Fig. 4C**), they showed no detectable binding to expired Novolog, despite these samples exhibiting a small but significant increase in fibril concentration by ThT assay. As with Humalog, this indicates that modest fibrillation arising from non-thermal degradation pathways may produce fibril structures that fall outside the antibodies’ recognition profiles.

Likewise, Basaglar showed no antibody binding in the fresh insulin controls but was readily detected after incubation at 37°C with agitation, which produced a high concentration of fibrils (**Fig. 4D**). Surprisingly, the antibodies also bound to Basaglar incubated at 65°C, even though this condition generated only a small, non-significant increase in fibril concentration by ThT assay. This could indicate that ThT does not efficiently recognize the fibrillar species formed in Basaglar at 65°C, or alternatively, that the antibodies preferentially detect fibril conformations induced by thermal stress, even when the overall fibril burden is low, highlighting their selective affinity for heat-induced structures.

## Discussion

We have shown that the visual quality-assurance tests recommended by insulin manufacturers fail to reliably detect insulin fibrils across different analogs and degradation conditions. Insulin degradation poses two distinct problems: the formation of fibrillar aggregates and the loss of functional insulin. Importantly, our results demonstrate that substantial fibril accumulation can occur without an accompanying reduction in insulin bioactivity, further limiting the ability of people with diabetes to recognize when their medication has been compromised. As a result, a PwD may unknowingly inject insulin containing high fibril burdens over prolonged periods, increasing the risk of long-term complications such as insulin-derived amyloidosis^18^.

Several consumer products marketed to people with diabetes attempt to address insulin instability by alerting users to temperature deviations, but these devices do not assess if degradation has actually occurred^4^. Our results demonstrate that this strategy is insufficient, as insulin degradation can arise from multiple environmental stressors beyond temperature alone, including UV exposure, air, and mechanical agitation (**Fig. 1B, Fig. S3**). Compounding this complexity, different insulin analogs exhibit distinct susceptibilities, with some analogs remaining stable under conditions that degrade others (**Fig. S3, Supplementary Data 1**). Furthermore, temperature monitoring alone fails to capture the synergistic effects of multiple stressors acting on insulin. For example, while Basaglar remains stable at 37°C under static conditions, the same temperature exposure leads to substantial fibrillation when agitation is introduced (**Fig. 1B, Fig. S3-4**). In addition, the extent of insulin degradation can be difficult to predict, as Basaglar’s degradation varied both within a single vial (**Fig. 2J**) and between different vials (**Fig. 2K**) when exposed to identical stressors. Taken together, these findings demonstrate that insulin integrity cannot be reliably inferred from temperature monitoring alone, underscoring the need for direct, product-level assessment of insulin degradation.

Given this unpredictability, we next explored whether antibody-based detection could provide a practical method to identify fibrils in degraded insulin. We demonstrate the feasibility of this approach by successfully detecting thermally degraded insulin analogs using a commercial fibril antibody kit^38^ (**Fig. 4**). Among the stressors examined, heat exposure produced the highest fibril concentrations (**Fig. 1B, Fig. S3**), suggesting that thermal degradation may represent a particularly significant risk for insulin instability. These experiments provide an initial validation of antibody-based detection as a potential strategy for assessing insulin quality. However, real-world insulin handling over weeks or months, often involving exposure to multiple stressors (**Fig. S4E-F**), may generate substantially greater fibril burdens and a wider diversity of fibrillar species than those measured here, reinforcing the need for point-of-care tests capable of detecting a broad spectrum of insulin degradation products.

Further studies should therefore focus on developing or testing antibodies capable of detecting fibrils generated through non-thermal degradation pathways, which our data indicate produce structurally distinct fibrillar conformations that fall outside the recognition profiles of the antibodies we tested (**Fig. 4**). In addition, future work should assess whether the commercial antibodies evaluated in this study can detect low concentrations of thermally formed insulin fibril proteins. For example, although exposure of insulin analogs to 37°C with agitation produced only gradual increases in degradation levels as measured by the ThT assay, the dot blot results suggest that antibody-based detection may have a lower effective limit of detection. This is supported by the Basaglar 65°C condition (**Fig. 4**), in which robust antibody binding was observed despite only a small, statistically insignificant increase in ThT fluorescence, suggesting that the antibodies may recognize early or structurally specific fibrillar intermediates.

Our results highlight a critical gap in current insulin quality assurance and establish antibody-based fibril protein detection as a promising path toward a point-of-use diagnostic. Such a tool would empower people with diabetes to better assess insulin integrity, reduce exposure to compromised medication, and mitigate the risk of long-term complications associated with injecting degraded insulin.

## Methods

### Insulin

Experiments were conducted with new, unopened insulin vials and pens of 100 unit/mL Humalog (Eli Lilly), Novolog (Novo Nordisk), and Basaglar (Eli Lilly). All insulin was stored in a Styrofoam box inside a 4°C fridge, keeping the insulin within temperature storage recommendations while shielding it from light^40^. Unless otherwise noted, insulin was extracted from its container using a needle no more than 28 days before the container’s first puncture.

### Insulin degradation procedures

For each degradation experiment, insulin was removed from its main insulin vial or insulin pen and added to three 2 mL microcentrifuge tubes (USA Scientific), aliquotting 100 μL per tube. Fresh, unaltered insulin was kept in a 4°C fridge (Danby) as a negative control. For the 37°C degradation trial, each insulin tube was kept in a heat block (Thermo Scientific) inside the top rack of a temperature-controlled incubator (Thermo Scientific MaxQ 6000) operating at 37°C. For the agitation degradation trial, each tube was placed in a shaking rack in the incubator at room temperature (around 23°C) at 220 RPM. The incubator’s heating element was turned off for the agitation trial. For the 37°C with agitation degradation trial, each tube was placed in a shaking rack in the 37°C incubator at 220 RPM. For the 65°C trial, each tube was placed in a heat block inside an incubator (Boekel Scientific) operating at 65°C, which can be the temperature in areas inside a hot car^41^. The temperature in each section of the incubators was verified with an external thermometer. For the -20°C trial, each tube was placed in a temperature block inside a -20°C freezer (Danby). Prior to sampling, it was noted if an insulin tube underwent a freeze-thaw cycle (**Fig. S3H**). For the air degradation trial, 500 μL of insulin was transferred to a 50 mL tube to maximize the air’s head space inside the tube. To avoid contamination and ensure continuous air exposure, the tube caps were removed and replaced with Parafilm (Sigma-Aldrich). The tubes were then kept inside a dark room at room temperature for the duration of the experiment. Instead of Parafilm, the experiment was repeated separately with either Breathe-Easy membranes (Sigma-Aldrich) or with the 50 mL cap on but unscrewed to limit evaporation, but these alternative methods had a negligible effect compared to the Parafilm air exposure trial (data not shown). For the UV degradation trial, 500 μL of insulin was transferred to a 25 mL glass tube to maximize UV exposure. The insulin was exposed at room temperature using a transilluminator box operating at 302 nm (UVP 2UV Transilluminator). The UV degradation trial was only run for 2 hours due to the rapid deterioration of insulin samples from the focused UV light.

### Absorbance assay

Insulin samples were aliquotted and exposed to various stressors for up to 4 days. The insulin was then assayed for absorbance at the end of experimentation using a spectrophotometer at 600 nm (Thermo Fisher NanoDrop 2000). Each measurement was performed in triplicate, using samples taken from independent microcentrifuge tubes. Representative images of the assayed insulin can be seen in **Fig. S1**. The average background of fresh insulin absorbance was minused from all samples before plotting.

### ThT fluorescence assay

ThT stock solutions were created from ThT salt (Sigma-Aldrich) diluted in autoclaved deionized water. Stocks were stored at 4°C while wrapped in aluminum foil for light protection. Insulin was aliquotted and exposed to various stressors for up to 4 days. The insulin was assayed for fibrillation at regular intervals using a plate reader (Tecan Infinite 200 Pro) with an excitation wavelength of 430 nm and an emission wavelength of 485 nm (**Supplemental Data 1**). Each measurement was performed in triplicate, using samples taken from independent microcentrifuge tubes, and then normalized by the fluorescence of fresh insulin. For this assay, insulin and Thioflavin T were used at the optimal concentrations of 297 μM and 16 μM, respectively (**Fig. S2**), and were mixed together with a pipette and incubated in the dark at room temperature for 10 minutes before measurement.

To determine the optimal concentrations for detection, Thioflavin T and the insulins were run in a gradient matrix, with the insulin either (a) exposed to a large stress known to cause high levels of fibril proteins or (b) unexposed to any stressor. Therefore, the concentrations producing the largest gain over background fluorescence were chosen for the Thioflavin T assay (**Fig. S2**).

### Mammalian cell culture

CHO INSR 1284 (ATCC CRL-3307) is a Chinese Hamster Ovary line that expresses insulin receptor B. These cells were maintained at 37°C in 5% CO2 in F-12 Glutamax media (Fisher Scientific #31765035),10% v/v FBS (Sigma #F4135), and 0.03% v/v of 50mg/mL Hygromycin (Sigma #10843555001), and passaged with accutase (Sigma #A6964).

### CHO insulin bioactivity assay

The researchers closely followed the 96-well plate insulin bioactivity assay protocol outlined by the FDA^34^. Briefly, insulin of various fibril protein levels were diluted and added to CHO cells that express the insulin receptor. When insulin binds its receptor, it triggers the auto-phosphorylation of tyrosine residues on the receptor. Insulin-induced auto-phosphorylation of the insulin receptor is determined as a read-out for insulin biological activity using a primary antibody (Sigma #05-321) that probes the phosphorylation cascade and a fluorescent secondary antibody for quantification (Fisher Scientific #A28175). A Hoechst stain (Fisher Scientific #62249) is then used to normalize the secondary antibody signal by cell number. While the original methodology compared commercial formulations to USP standards, we compared stressor-exposed commercial formulations to unaltered commercial formulations.

We selected environmental stressors that produced undetectable, low, medium, and high levels of fibril proteins in one or more of the analogs. All three insulin analogs were tested under five core conditions: 37°C with agitation, 65°C, freeze–thaw cycles at -20°C, expiration, and ideal storage at 4°C. To more precisely characterize degradation at their most sensitive conditions, Humalog and Novolog were incubated at 65°C for 6 to 48 hours, and Basaglar was incubated at 37°C with agitation for 3 to 7 days. We selected 48-hour incubations for Humalog and Novolog at 37°C with agitation, and for Basaglar at 65°C, because fibril protein levels showed only minimal increases beyond these timepoints (**Fig. S3**). -20°C trials consisted of three freeze-thaw cycles over two days. In addition, we included insulin samples past their expiration dates due to its relevancy for PwD^17^. For the “Expired” conditions, Novolog was used from an unpunctured insulin vial that expired in 2020 that was donated by a PwD, while Humalog and Basaglar were from pens first punctured three and four months earlier by the researchers, respectively. The full list of insulin tested in the CHO assay is provided in **Supplemental Data 2**.

Two to five replicates of each experimental condition were tested on each CHO plate. Each plate contained wells with fresh, unaltered insulin added to the CHO cells as a positive control. Because some stress conditions markedly reduced insulin bioactivity resulting in flat response curves across dilutions, the EC_50_ value – which quantifies the insulin concentration causing a 50% bioactivity response – was not an appropriate measure of potency. Instead, bioactivity was quantified using the area under the curve (AUC) of two to three representative insulin dilutions that best reflected differences in bioactivity and were near the EC_50_ value of fresh insulin (**Fig. S5**).

To quantify insulin bioactivity, the fluorescence intensity of the secondary antibody signal was first divided by the Hoechst stain signal to normalize for the number of CHO cells per well. Then, the AUC of insulin bioactivity across dilutions was calculated for each experimental condition. Next, the mean AUC from PBS-treated wells was subtracted from each experimental condition. However, for a Novolog plate that lacked PBS-treated wells, the lowest degraded insulin dilution was used for background subtraction (∼10^-5^ µM), which had minimal bioactivity compared to the dilutions used for analysis (above ∼10^-2^ µM) (**Fig. S5B**) and had similar AUC values as PBS-treated wells (**Fig. S7A**). Removal of this plate yielded qualitatively similar results (**Fig. S7B**). Finally, each experimental AUC was normalized to the AUC of the fresh insulin replicates (**Supplemental Data 2**). The normalized CHO bioactivity readings were then compared to fibril protein concentration, which was determined by ThT assay.

### Protein secondary structure determination by circular dichroism

Insulin was aliquotted to 2 mL microcentrifuge tubes and exposed to the stressor that formed the highest concentration of fibril protein as determined by ThT assay. Therefore, Humalog and Novolog were exposed to 65°C, and Basaglar was exposed to 37°C with agitation (**Fig. 1B**). Each insulin produced high, medium, and undetectable fibril protein concentrations under their stress condition, with 3 samples per concentration, for a total of 9 samples per insulin. The insulin samples were then diluted 1:167 from their original formulation in autoclaved deionized water and analyzed in a CD spectrophotometer (JASCO J-815) using a 5 mm pathlength quartz cuvette (Labomed). This protein concentration kept the circular dichroism HT voltage below ∼600 V at the farthest UV wavelengths tested while keeping the magnitude of the spectra minima of fresh insulin near 10 mdeg. Before analyzing experimental samples in the CD, a baseline spectra of autoclaved deionized water was taken, which was subtracted from the experimental spectra. The insulin samples were analyzed using the following CD settings for both experimental and water baseline measurements: a digital integration time of 2 seconds, band width of 1 nm, data pitch of 0.1 nm, scanning speed of 50 nm/min, continuous scanning mode, and standard sensitivity from 250 nm to 200 nm for 10 accumulations.

### Antibody affinity to insulin fibril proteins using dot blot

Insulin was aliquotted to 2 mL microcentrifuge tubes and then degraded in two different ways per insulin type. The different degradation types were chosen to see if degradation type would affect the antibody affinity, and because they form differing fibril concentrations in their respective insulin analog (**Fig. 4A**). Specifically, the insulin analog samples were exposed to 65°C for 48 hours, three freeze-thaw cycles over two days at -20°C, expired storage, 37°C with agitation for 10 days, or ideal storage conditions at 4°C.

The insulin was then assayed for degradation using the ThT assay. A commercially-available antibody panel that recognizes general epitopes of fibrils’ secondary structure (Abcam)^38,39^ was then assayed with the insulin samples.

For the dot blot protocol, insulin was first diluted to 3 concentrations in TBS: 1 mg/mL, 0.1 mg/mL, and 0.01 mg/mL. The fourth dilution in the Novolog mOC87 blot was 0.001 mg/mL (**Fig. S8B**). Then, 3 μL of the insulin sample was spotted on 0.2 μm nitrocellulose membrane (Sigma-Aldrich) and allowed to dry. The blot was gently agitated for 1 hr in TBST+2% BSA blocking solution (Sigma-Aldrich) at room temperature. Next, the sample was placed in a 1:8,000 primary antibody blocking solution overnight at 4°C while gently agitated. The next day, the primary antibody solution was poured into a 15 mL tube (Falcon), and the tube was placed into a 4°C refrigerator for future use. After three washes for 5 minutes each with TBST at room temperature, the blot was gently agitated in 1:20,000 IRDye® 800CW Goat anti-Rabbit IgG Secondary Antibody (LICORBio) blocking solution for 1 hr at room temperature. To mitigate light exposure before analysis, the blot carrier was covered in aluminum foil during the secondary antibody blocking step and all subsequent steps. After three washes for 5 minutes each in TBST, TBS was added after the final wash. The blot was then removed from its carrier and imaged (LI-COR Odyssey CLx dual-color imager). For these dot blot assays, TBS was the negative control, and Aβ42 was the positive control. Although the antibody panel had variable affinity to Aβ42 in past studies^39^, the panel had sufficient affinity to Aβ42 at our tested protein and antibody dilutions. In all blots, Aβ42 was diluted 1:100 unless otherwise noted.

A custom Matlab script was used to change the brightness of all images, change all images to a white background with black dot, and crop the dot blots into an image matrix, which created **Fig. 4C-E**.

### Significance in pairwise comparisons

The assumption of normal distributions in the data was checked using the Anderson-Darling test before proceeding with further statistical testing. For pairwise comparisons between normally distributed data, significance was determined using a two-way t-test with an α value of 0.05. For multiple comparisons, ANOVA was performed using the Tukey-Kramer correction when checking each pairwise comparison and the Dunnett correction when checking each experimental condition compared only to a control. The asterisks denote ***** p < 10^-5^, **** p < 10^-4^, *** p < 10^-3^, ** p < 10^-2^, and * p < 5*10^-2^.

## Data availability

The data that support the findings of this study are openly available. All data and code generated in the study are included in the SI and at our GitHub page: https://github.com/davidritz/insulin. Further inquiries can be directed to the corresponding author.

## Ethical considerations

This article does not contain any studies with human or animal participants.

## Consent for publication

Not applicable.

## Consent to participate

Not applicable.

## Acknowledgements.

We thank Aaron McKenna for the careful reading and valuable suggestions to our manuscript. We thank Thao Huyhn for important discussions on our experimental procedures. We thank Alex Fu for instruction on dot blot protocols, Paul DeFino and Pepper Pennington for important guidance on the CD experiments, and Dean Madden for use of his lab’s quartz cuvette for CD experimentation. We thank Jen Bomberger and TY Chang for allowing the storage of our CHO cells in their labs. Equipment and tools from BioMT were used, which is supported through NIH NIGMS grant P20-GM113132.

## Author contributions

D.R. designed the study. D.R. performed the ThT, CD, and absorbance experiments. D.R. prepared the insulin samples, and E.L.S. performed the tissue culture assay for the CHO experiments. D.R. analyzed the data. E.L.S. and D.S. provided important edits on the manuscript. D.R. provided lab materials. D.S. provided oversight and lab space. D.R. wrote the manuscript with input from all authors.

## Statements and Declarations

Not applicable.

## Supplemental Files Information

**Ritz_jdr_supp2.docx:** Provides supplemental figures S1-S8 that support the main text.

**Supplemental Data 1.xlsx:** Table of ThT kinetic data and absorbance data for each insulin type and degradation type.

**Supplemental Data 2.xlsx:** Table of CHO bioactivity and ThT data for each insulin type and degradation type.

