## Supplemental Information for "Towards the development of an insulin degradation test"

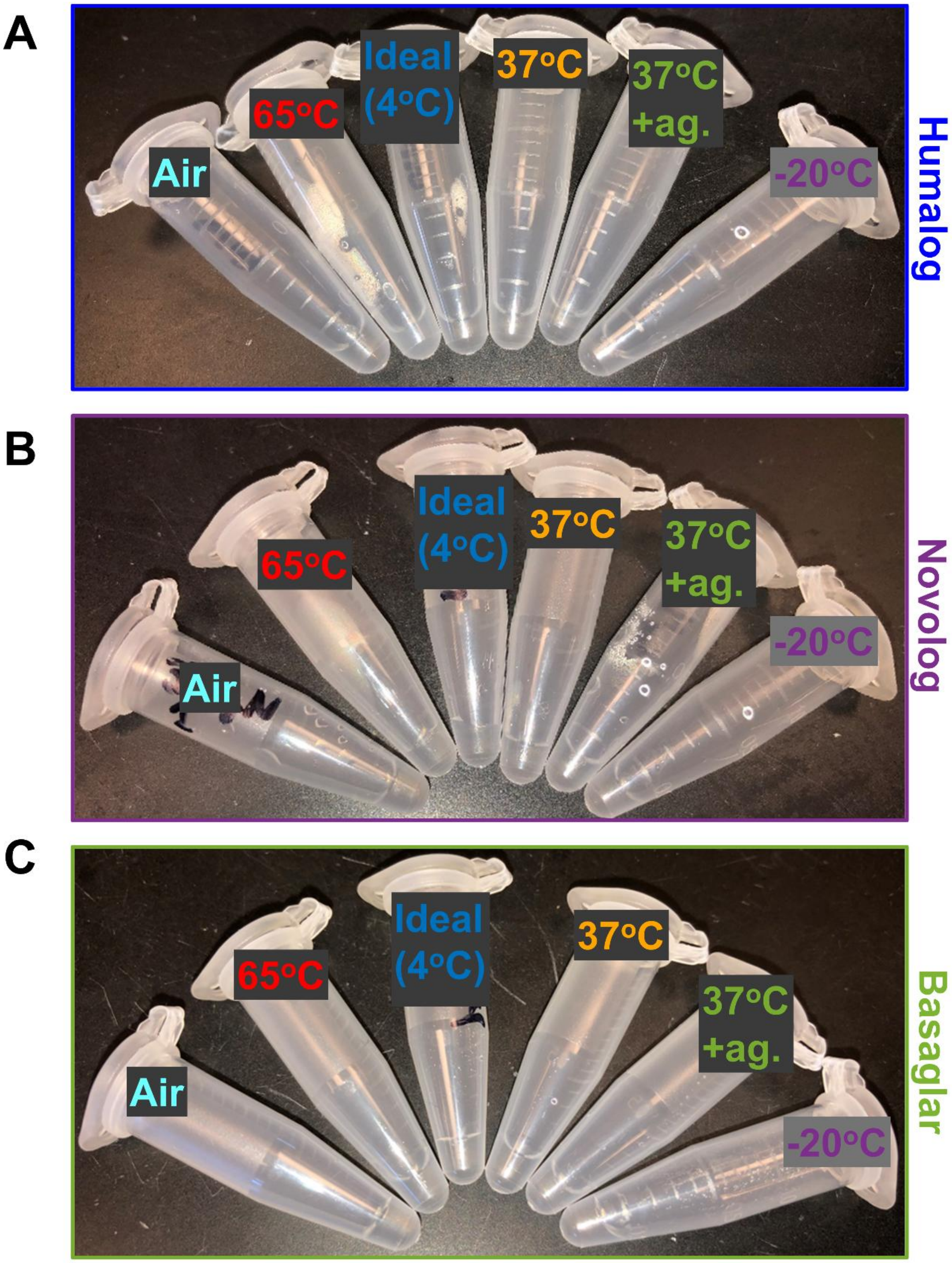


**Figure S1: Insulin rarely turns cloudy from acute degradations.** Representative images of **(A)** Humalog, **(B)** Novolog, and **(C)** Basaglar insulin samples in microcentrifuge tubes after 96 hours of degradation in each of the labeled stress conditions. UV and agitation samples are not shown.

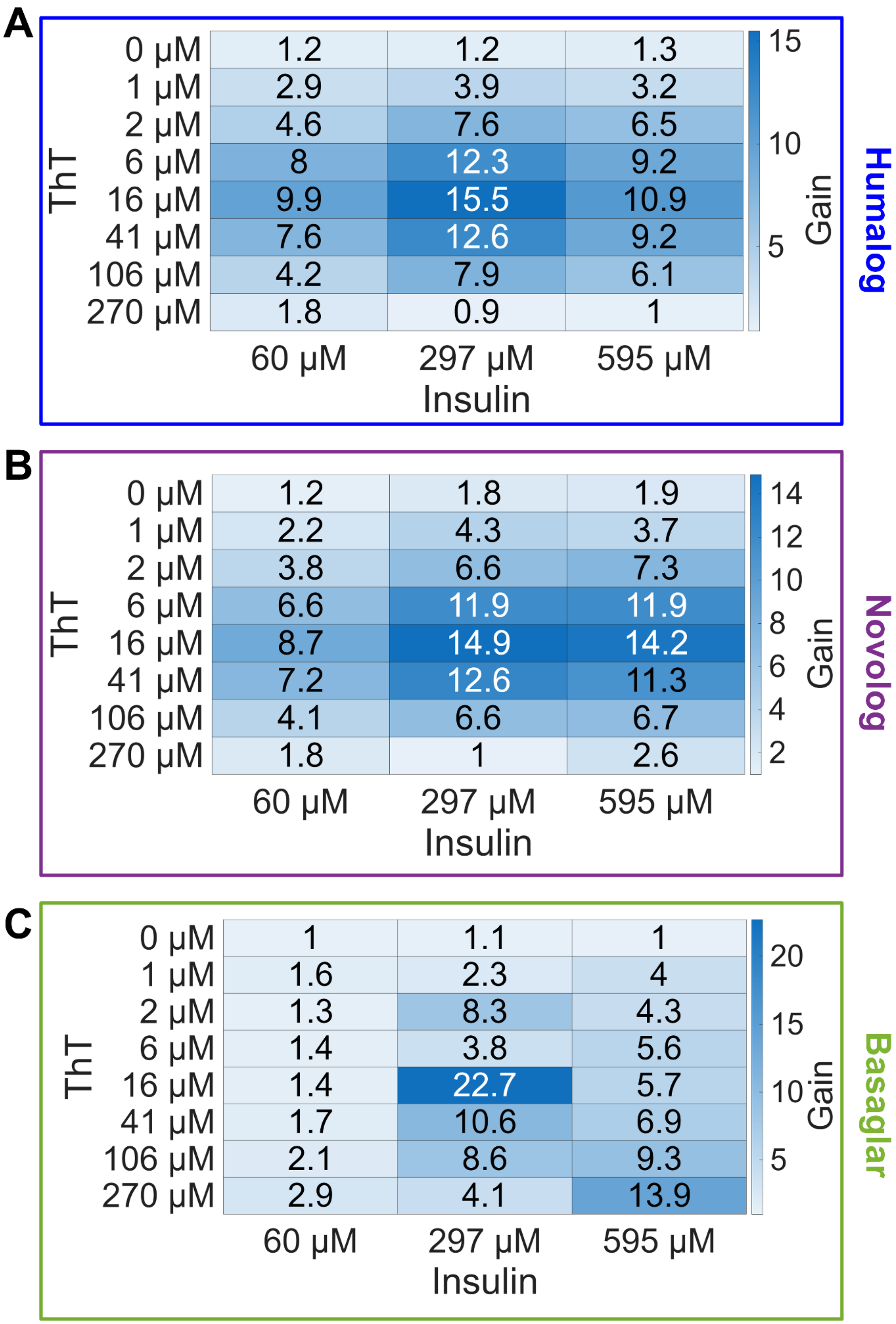
A
Humalog
ThT
0 μM 1.2 1.2 1.3
1 μM 2.9 3.9 3.2
2 μM 4.6 7.6 6.5
6 μM 8 12.3 9.2
16 μM 9.9 15.5 10.9
41 μM 7.6 12.6 9.2
106 μM 4.2 7.9 6.1
270 μM 1.8 0.9 1
60 μM 297 μM 595 μM
Insulin
Gain
15 10 5
B
Novolog
ThT
0 μM 1.2 1.8 1.9
1 μM 2.2 4.3 3.7
2 μM 3.8 6.6 7.3
6 μM 6.6 11.9 11.9
16 μM 8.7 14.9 14.2
41 μM 7.2 12.6 11.3
106 μM 4.1 6.6 6.7
270 μM 1.8 1 2.6
60 μM 297 μM 595 μM
Insulin
Gain
14 12 10 8 6 4 2
C
Basaglar
ThT
0 μM 1 1.1 1
1 μM 1.6 2.3 4
2 μM 1.3 8.3 4.3
6 μM 1.4 3.8 5.6
16 μM 1.4 22.7 5.7
41 μM 1.7 10.6 6.9
106 μM 2.1 8.6 9.3
270 μM 2.9 4.1 13.9
60 μM 297 μM 595 μM
Insulin
Gain
20 15 10 5

**Figure S2: Optimal ThT and insulin concentrations used during ThT assay.** Heat maps of ThT fluorescence gain for **(A)** Humalog, **(B)** Novolog, and **(C)** Basaglar degraded insulin samples relative to fresh insulin samples. To generate the degraded samples, Humalog and Novolog were degraded at 65$^{o}$C for 24 hours and Basaglar was degraded at 37$^{o}$C with agitation for 4 days. Because of these results, all ThT assays used an insulin concentration of 297 μM and a ThT concentration of 16 μM.

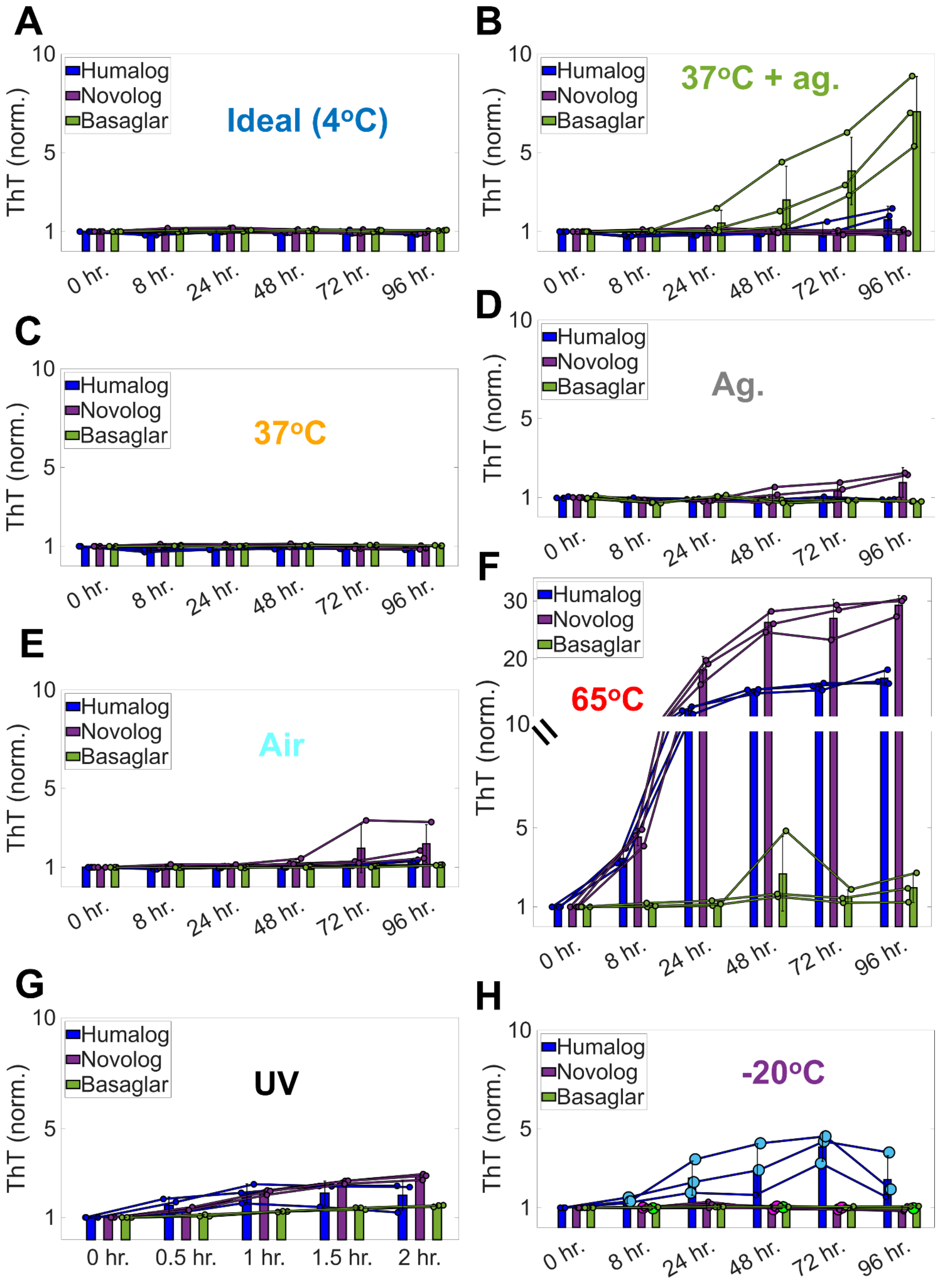

A
Ideal (4oC)
B
37oC + ag.
C
37oC
D
Ag.
E
Air
F
65oC
G
UV
H
-20oC
Humalog
Novolog
Basaglar
ThT (norm.)
0 hr.
8 hr.
24 hr.
48 hr.
72 hr.
96 hr.
0.5 hr.
1 hr.
1.5 hr.
2 hr.

**Figure S3: Kinetics of fibril protein formation differ by stressor type and insulin type.** The kinetics of fibril protein formation was tested for three aliquots of each insulin type when continuously exposed to: **(A)** ideal conditions at 4$^{o}$C, **(B)** 37$^{o}$C with agitation, **(C)** 37$^{o}$C, **(D)** agitation, **(E)** air, **(F)** 65$^{o}$C, **(G)** 302 nm UV light, and **(H)** -20$^{o}$C. The large dots in panel H correspond to when the insulin went through a freeze-thaw cycle before sampling for the ThT assay. This was denoted because the insulin did not always freeze when in the -20$^{o}$C freezer. The ThT fluorescence at the end of each experiment is displayed in **Fig. 1B**.

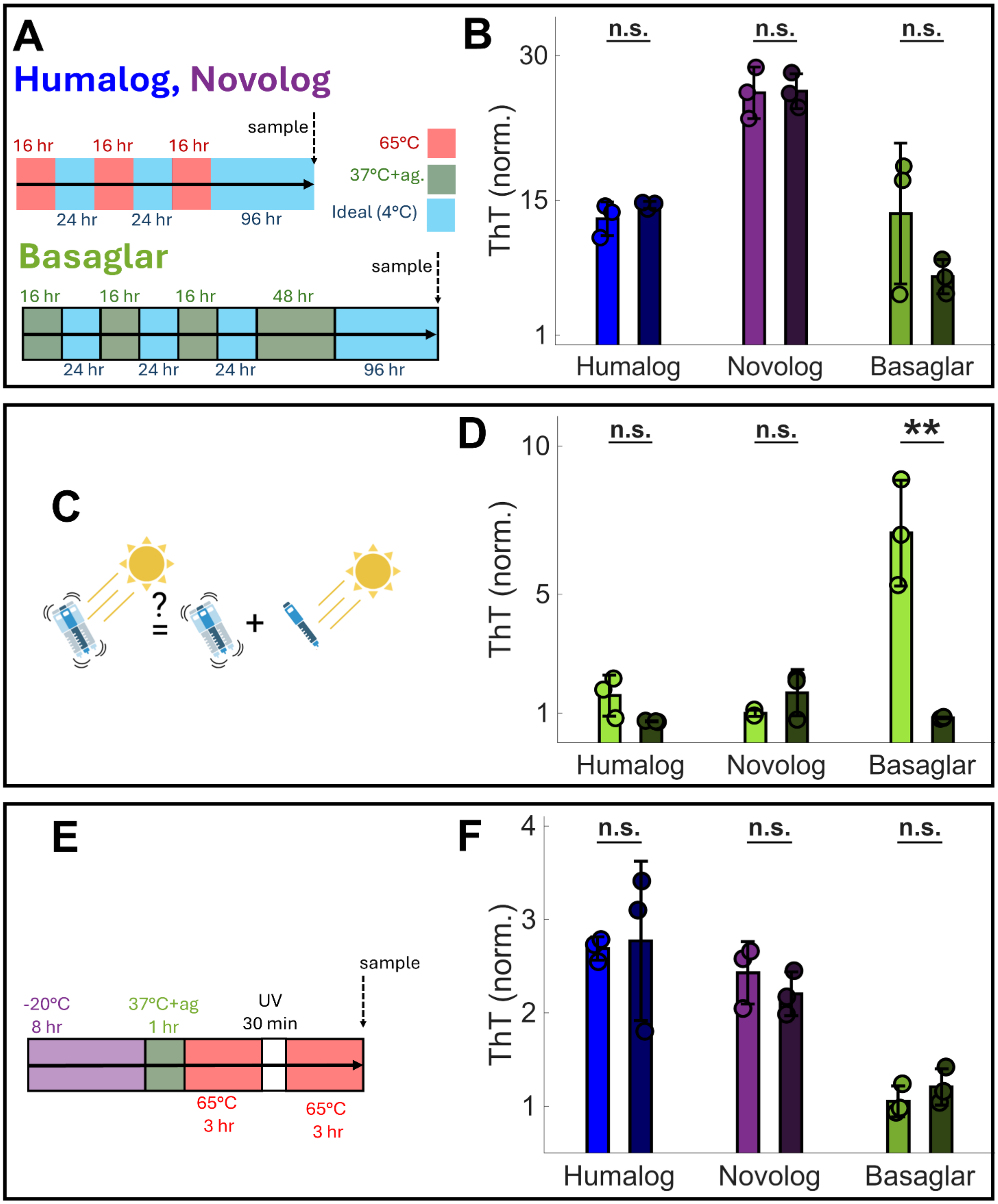

A
Humalog, Novolog
16 hr
16 hr
16 hr
sample
24 hr
24 hr
96 hr
65°C
37°C+ag.
Ideal (4°C)
Basaglar
sample
16 hr
16 hr
16 hr
48 hr
24 hr
24 hr
24 hr
96 hr
B
n.s.
n.s.
n.s.
30
15
1
ThT (norm.)
Humalog
Novolog
Basaglar
C
?
=
+
D
n.s.
n.s.
**
10
5
1
ThT (norm.)
Humalog
Novolog
Basaglar
E
sample
-20°C
8 hr
37°C+ag
1 hr
UV
30 min
65°C
3 hr
65°C
3 hr
F
n.s.
n.s.
n.s.
4
3
2
1
ThT (norm.)
Humalog
Novolog
Basaglar

**Figure S4: Insulin degradation is generally additive and irreversible, but Basaglar 37$^{o}$C with agitation is not additive. (A-B)** To assess the reversibility of degradation, each insulin type was exposed to rounds of a degradative stressor and was placed in ideal conditions at the end of each round. The lighter bar represents the observed degradation at the end of experimentation, while the dark bar represents the expected degradation from an additive model, calculated by adding the degradation of singular stressors. A sign of degradation reversibility would be a decrease in fibril protein formation relative to the additive model. **(C-D)** Insulin was agitated while at 37$^{o}$C, and its degradation was compared to the same insulin separately agitated and then exposed to 37$^{o}$C. The lighter bar represents the observed degradation when insulin is exposed to both stressors simultaneously, while the darker bar represents the expected degradation from an additive model, calculated by adding the degradation of singular stressors **(E-F)** Insulin was exposed to rounds of freezing temperatures, hot temperatures, UV light, and agitation, denoted as the "Bad Beach Day" trial (BBD) in **Supplemental Data 1**. The lighter bar represents the observed degradation at the end of experimentation, while the darker bar represents the expected degradation from an additive model, calculated by adding the degradation of singular stressors. For each experiment, 3 replicates were used for each trial. In each panel, the observed degradation was compared to the expected degradation by a two-way t-test. The statistics notation denote ***** $p < 10^{-5}$, **** $p < 10^{-4}$, *** $p < 10^{-3}$, ** $p < 10^{-2}$, and * $p < 5*10^{-2}$, and n.s. $> 5*10^{-2}$.

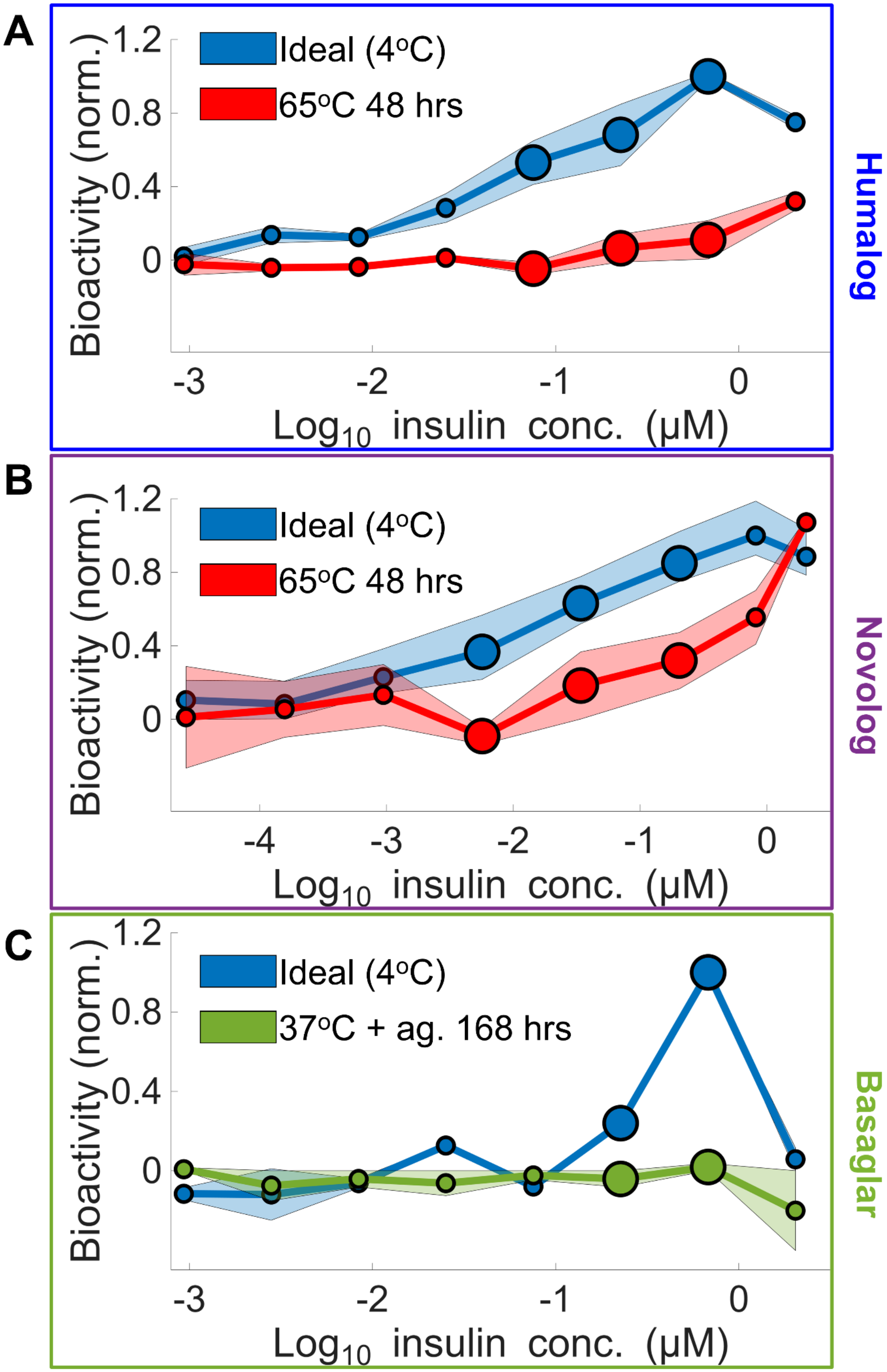

A
Ideal (4°C)
65°C 48 hrs
Bioactivity (norm.)
1.2
0.8
0.4
0
-3
-2
-1
0
Log10 insulin conc. (μM)
Humalog
B
Ideal (4°C)
65°C 48 hrs
Bioactivity (norm.)
1.2
0.8
0.4
0
-4
-3
-2
-1
0
Log10 insulin conc. (μM)
Novolog
C
Ideal (4°C)
37°C + ag. 168 hrs
Bioactivity (norm.)
1.2
0.8
0.4
0
-3
-2
-1
0
Log10 insulin conc. (μM)
Basaglar

**Figure S5: CHO assay reveals differences in bioactivity between fresh and degraded insulin at certain insulin dilutions.** Fresh insulin and degraded insulin's CHO bioactivity was compared for **(A)** Humalog, **(B)** Novolog, and **(C)** Basaglar. The large dots represent insulin concentrations that show large separations in bioactivity between fresh insulin and fibril-protein-rich insulin and are near the $EC_{50}$ value of fresh insulin, which is the insulin concentration causing a 50% bioactivity response. These concentrations were used for further analysis in the CHO bioactivity assay shown in **Fig. 2**. The shading around each line graph represents the maximum and minimum of two replicates.

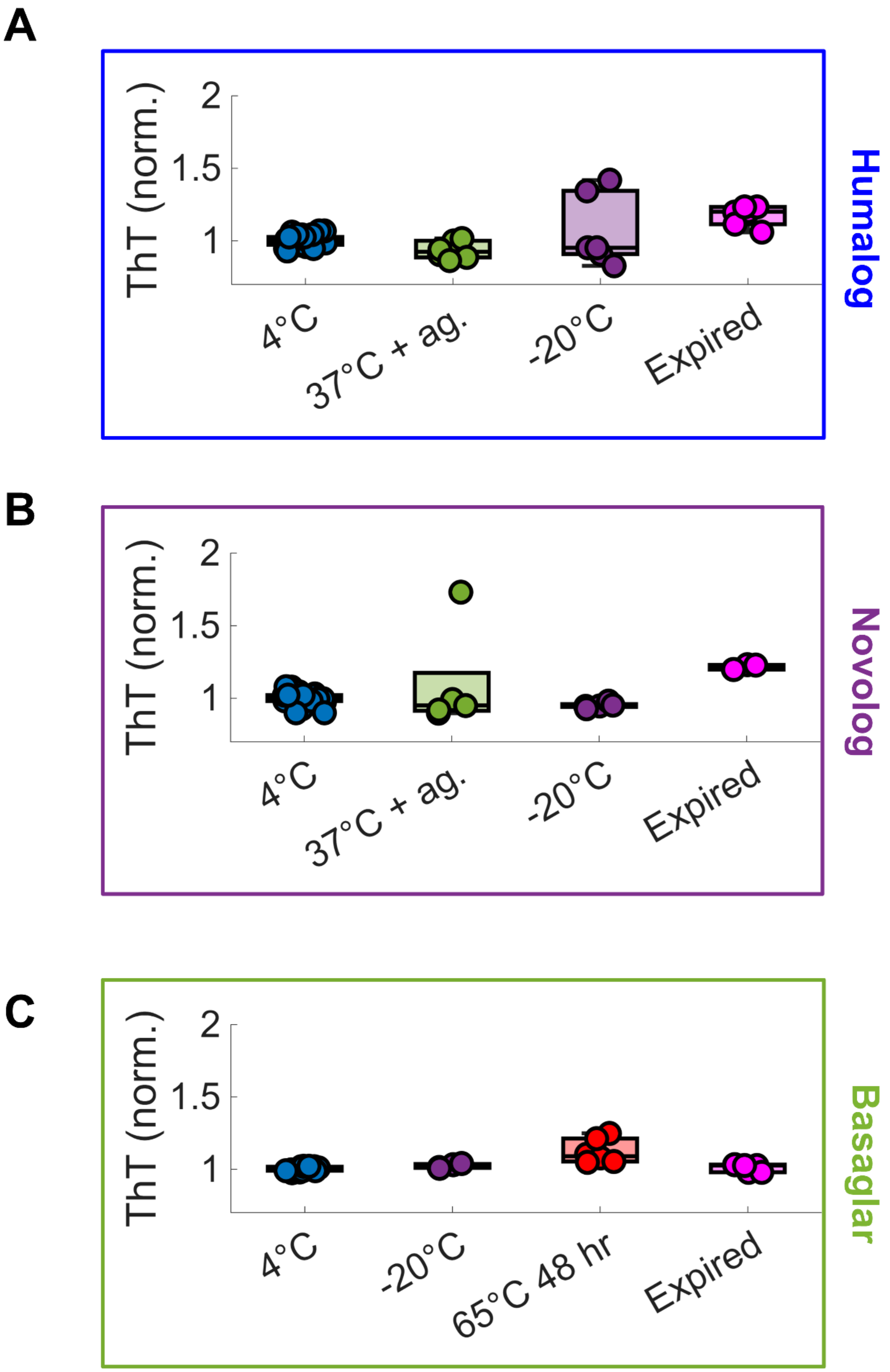

A
ThT (norm.)
2
1.5
1
4°C
37°C + ag.
-20°C
Expired
Humalog
B
ThT (norm.)
2
1.5
1
4°C
37°C + ag.
-20°C
Expired
Novolog
C
ThT (norm.)
2
1.5
1
4°C
-20°C
65°C 48 hr
Expired
Basaglar

**Figure S6: Stressors can produce small changes to insulin fibril protein concentration.** During the CHO experiment, stressors for **(A)** Humalog, **(B)** Novolog, and **(C)** Basaglar were chosen that produce small or no changes to ThT fluorescence (**Fig. S3**). All trials were compared to their respective insulin analog at 4$^{o}$C using ANOVA with Dunnett correction. This figure was created by rescaling **Fig. 2C**, **Fig. 2F**, and **Fig. 2I**.

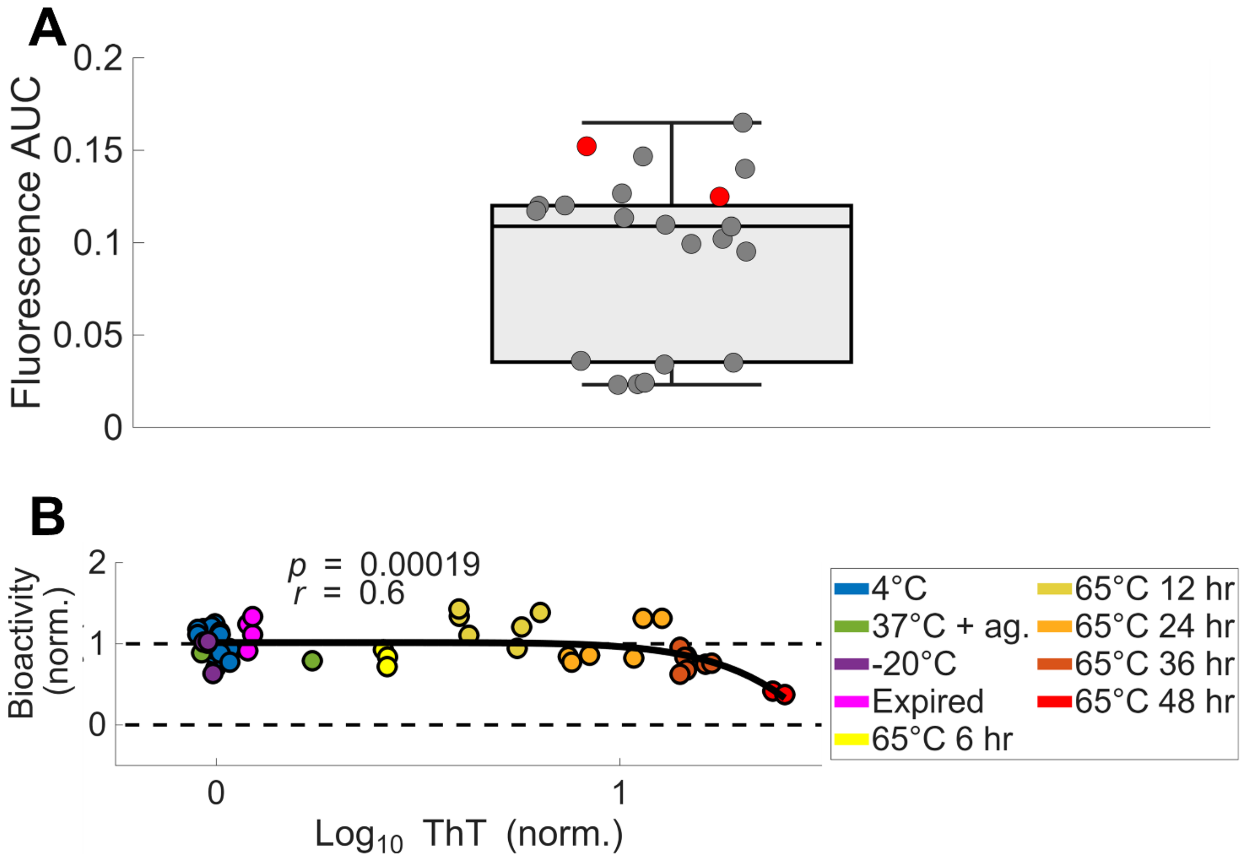


**Figure S7: Highly degraded and diluted Novolog insulin is an appropriate background measurement. (A)** Boxplot of AUC values across all plates using PBS-treated wells as background (gray), with individual PBS measurements shown as gray points. Overlaid are the AUC values using the lowest dilutions of Novolog incubated at 65°C for 48 h (red, **Fig. S5B**), exhibiting similarly low signals. Therefore, these samples serve as surrogate background measurements for the Novolog CHO plate lacking PBS-treated CHO cells. **(B)** CHO bioactivity versus fibril protein concentration for Novolog with the surrogate-background plate removed. All CHO plates are included in **Fig. 2D**, showing only minor differences with **Fig. S7B**. This confirms that use of the surrogate background minimally affects the results.

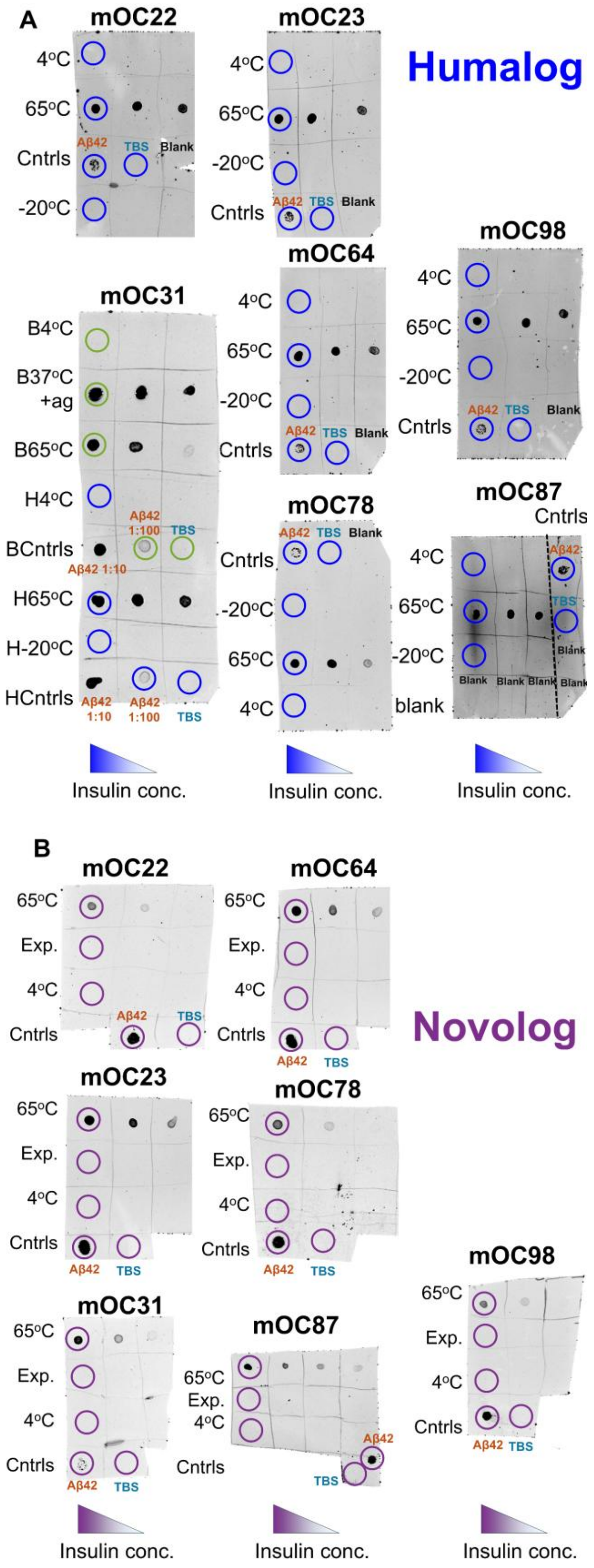
A
Humalog
mOC22
4oC
65oC
Aβ42
TBS
Blank
Cntrls
-20oC
mOC23
4oC
65oC
-20oC
Aβ42 TBS Blank
Cntrls
mOC31
B4oC
B37oC +ag
B65oC
H4oC
Aβ42 1:100
TBS
BCntrls
Aβ42 1:10
H65oC
H-20oC
HCntrls
Aβ42 1:10
Aβ42 1:100
TBS
mOC64
4oC
65oC
-20oC
Aβ42 TBS Blank
Cntrls
mOC78
Aβ42 TBS Blank
Cntrls
-20oC
65oC
4oC
mOC98
4oC
65oC
-20oC
Aβ42 TBS Blank
Cntrls
mOC87
Cntrls
Aβ42
TBS
Blank
4oC
65oC
-20oC
Blank Blank Blank Blank
blank
Insulin conc.
Insulin conc.
Insulin conc.
B
Novolog
mOC22
65oC
Exp.
4oC
Aβ42
TBS
Cntrls
mOC64
65oC
Exp.
4oC
Cntrls
Aβ42
TBS
mOC23
65oC
Exp.
4oC
Cntrls
Aβ42
TBS
mOC78
65oC
Exp.
4oC
Cntrls
Aβ42
TBS
mOC31
65oC
Exp.
4oC
Cntrls
Aβ42
TBS
mOC87
65oC
Exp.
4oC
Aβ42
TBS
Cntrls
mOC98
65oC
Exp.
4oC
Cntrls
Aβ42
TBS
Insulin conc.
Insulin conc.
Insulin conc.

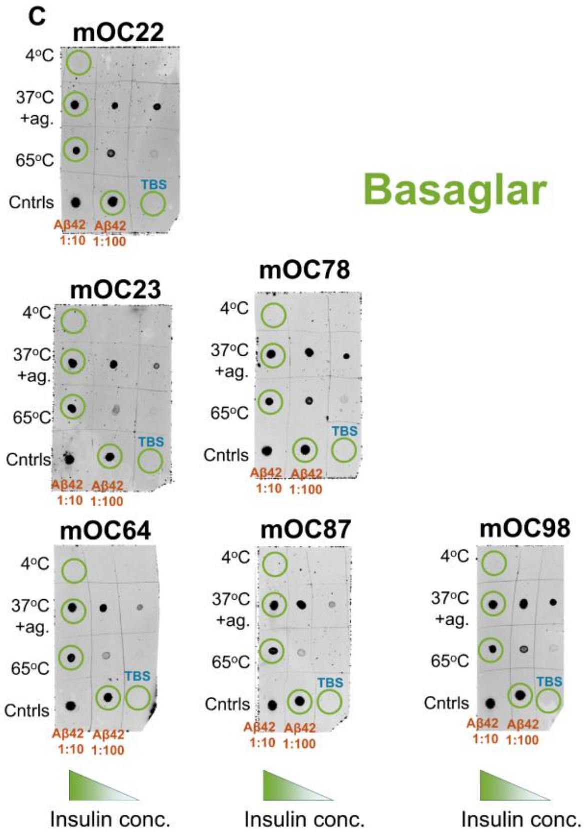


**Figure S8: Antibodies differ in their affinity to fibril proteins by degradation type.** **(A)** Humalog, **(B)** Novolog, and **(C)** Basaglar dot blots were conducted with antibodies that recognize general epitopes of fibril's secondary structure and with insulin that was degraded in three different ways per insulin type. Three dilutions for each insulin were tested: 1 mg/mL, 0.1 mg/mL, and 0.01 mg/mL. The fourth dilution in Novolog mOC87 is 0.001 mg/mL. Basaglar and Humalog blots were analyzed together on the same membrane for mOC31 in panel **A**. The circles denote which dots were used to construct **Fig. 4C-E**, which were the highest insulin concentrations tested. Aβ42 was the positive control, and TBS was the negative control. Aβ42 was diluted 1:100 in each blot unless otherwise noted.
